## Supplemental Information for "Context-Seq: CRISPR-Cas9 Targeted Nanopore Sequencing for Transmission Dynamics of Antimicrobial Resistance"

### Supporting Information

**List of Tables**


| List of Tables | Page |
|---|---|
| Supplementary Table 1: Taqman Array Card assays | 3 |
| Supplementary Table 2: Samples selected for Context-Seq | 4 |
| Supplementary Table 3: Candidate guide RNAs for CTX-M and TEM | 5 |
| Supplementary Table 4: CTX-M alleles captured with CTX-M guides used in this study | 5 |
| Supplementary Table 5: TEM alleles captured with TEM guides used in this study | 6 |
| Supplementary Table 6: MinION flow cell ID numbers and SRA sample numbers | 7 |
| Supplementary Table 7: Sequencing summary for samples used in this work | 7 |


**List of Figures**


| List of Figures | Page |
|---|---|
| Supplementary Figure 1: Percentage of sample positive for ARGs and pathogen genes | 4 |
| Supplementary Figure 2: Comparison of TEM guide RNAs for enrichment in model system. | 5 |

**Commercial Materials**
Powersoil Pro kit (Qiagen, 47016), Taqman array cards (Thermo Fisher Scientific, custom order), TaqPath (Applied Biosystems, A15299), crRNAs (IDT, custom), tracrRNAs (IDT, 1072532), duplex buffer (IDT, 11-01-03-01), CutSmart (NEB, B7204), HiFi Cas9 Nuclease V3 (IDT, 1081060), QuickCIP (NEB, M0525S), Thermolabile Proteinase K (NEB, P8111S), dATP (Invitrogen, 10216018), Taq polymerase (NEB, M0273S), Quick Ligation Kit (NEB, M2200S), Adapter Ligation Kit (ONT, SQK-LSK110) AMpure Beads (Beckman Coulter, A63880).

**Experimental Methods**
***Standard Preparation***
*E. coli* were isolated from the stool of children participating in a 2016-2019 cohort study of enteric infections in Lima, Peru (NIH R01AI108695-01A1).[1] Lactose fermenting and non-fermenting colonies were first isolated on MacConkey agar, then sub-plated on Chromagar ESBL to identify presumptive ESBL-producers. ESBL production was phenotypically confirmed using the double disk diffusion assay. The isolate containing CTX-M-55 and TEM-1 used for validation in this work was isolated from a 6-month-old male. For validation experiments, the isolate was plated on MacConkey agar and incubated at 37°C overnight. An individual colony was selected and cultured in 5 mL of LB overnight at 37°C. Media was pelleted and DNA was extracted using Qiagen's Powersoil Pro kit according to the manufacturer's instructions. DNA was quantified on a Qubit fluorometer.

***TaqMan Array Card***
Samples were pre-amplified with a pool of primers for all targets. A 10 µL reaction was set up with 2.5 µL of TaqPath MM (4x), 2.5 µL of PreAmp Pool, and 5 µL of DNA. Samples were run according to the following thermal cycling conditions: 2 min at 25°C, 1 min at 95°C, 15 cycles of 15 seconds at 95°C and 2 min at 60°C, followed by 10 min at 99°C. Pre-amplified samples were then diluted 20X in nuclease free water. Samples were amplified in a TaqMan Array Card in 100 µL reaction volume composed of 50 µL Taqman Fast Advanced Master Mix, 30 µL nuclease free water, and 20 µL diluted sample. Samples were run according to the following cycling conditions: 2 min at 50°C, 10 min at 92°C, 40 cycles of 95°C for 1 second and 60°C for 20 seconds.

**Supplementary Table 1:** qPCR assays used on a Taqman Array Card for 14 antibiotic resistance and 8 pathogen genes.

| Target | **Forward primer** (5'-3') | **Reverse primer** (5'-3') | **Probe** (5'-3') | Ref |
|---|---|---|---|---|
| TEM | CACTATTCTCAGAATGACTTGGT | TGCATAATTCTCTTACTGTCATG | CCAGTCACAGAAAAGCATCTTACGG | [2] |
| TEM-238S | GCTGGCTGGTTTATTGCTGATA | GCCCCAGTGCTGCAATG | ATCTGGAGCCAGTGAG | [3] |
| qnrS | TTGCTCAGCMTTTATTWCWGGATGT | CAGCGATTTTCAWACARCTCACA | TATGCCAATATGGAGMGGGT | [3] |
| sul1 | CGTGCTGTCGAACCTTCAAA | CCTTTACAGGAAGGCCAACG | AGAAGGATTTCCGCGACAC | [3] |
| ermB | CGTACCTTGGATATTCACCG | GTAAACAGTTGACGATATTCTCG | TGCACACTCAAGTCTCGATTCAGC | [3] |
| mcr-1 | GATCGCTGTCGTGCTCTTTG | ACCGCGCCCATGATTAATAG | CGATGCTACTGATCACCACG | [3] |
| CTX-M-1 | CCGTCACGCTGTTRTTAGGA | AATGCCACMCCCAGYCKKCC | CAGCAAAAACTTGCCGRATT | [3] |
| CTX-M-9 | GCTTTATGCGCAGACGARTG | ATCACCGCGATAAAGCACCT | TCGATACCRMAGATAATACGC | [3] |
| tetA | CGACGGCACAGGCTACATC | CCTGGACAACATTGCTTGCA | TGGCGTTCCCGATCAT | [3] |
| NDM | ATATCACCGTTGGGATCGAC | TAGTGCTCAGTGTCGGCATC | AAGGACAGCAAGGCCAAGTCG | [3] |
| OXA-48 like | GCAAAGGAATGGCAAGAAAA | CACAACTACGCCCTGTGATTT | AGTTGGAATGCTCACTTTACTG | [3] |
| OXA-10 | CAGAGAAGTTGGCGAAGTAAGA | ATTCTGGTTGGAAGGCCAG | CCTATGGCAACCAGAATATCAGTGGTGG | This study |
| SHV 238-240SE-SK | GTTGATCCGYTCCGTGCT | GCTTTGTTATKCGGGCCAAG | CGGAGCTAGCRARC | [3] |
| SHV | TCCCATGATGAGCACCTTTAAA | TCCTGCTGGCGATAGTGGAT | TGCCGGTGACGAACAGCTGGAG | [3] |
| C. jejuni | CTTGCGGTCATGATGGACATAC | AGCACCACCCAAACCCTCTTCA | TGCTTGCTGCAAAGTATT | [4] |
| C. coli | AAACCAAAGCTTATCGTGTGC | AGTCCAGCAATGTGTGCAAT | TAAGCTCCAACTTCATCCG | [4] |
| S. enteritidis | CTTTCTCAGATTCAGGGAGTATATCA | TGAACTACGTTCGTTCTTCTGGT | ATCAGCCTGTTGTCTGCTCACCAT | [5] |
| S. enterica | TCGGGCAATTCGTTATTGG | GATAAACTGGACCACGGTGACA | AAGACAACAAAACCCACCGC | [6] |
| Shigella | CCTTTTCCGCGTTCCTTGA | CGGAATCCGGAGGTATTGC | CGCCTTTCCGATACCGTCTCTGCA | [4] |
| STEC | CCACATCGGTGTCTGTTATTAACC | GGTCAAAACGCGCCTGATAG | TTGCTGTGGATATACGAGG | [4] |
| ETEC | TTCCCACCGGATCACCAA | CAACCTTGTGGTGCATGATGA | CTTGGAGAGAAGAACCCT | [4] |
| EPEC | CATTGATCAGGATTTTTCTGGTGATA | CTCATGCGGAAATAGCCGTTA | ATACTGGCGAGACTATTTCAA | [4] |

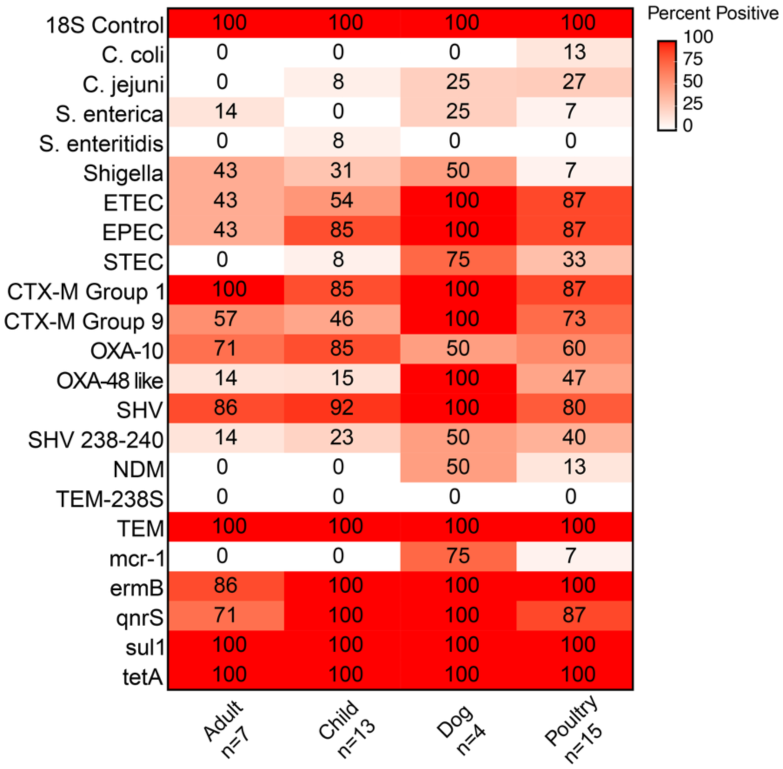


**Supplementary Figure 1**: Percentage of samples positive for 14 ARGs and 8 pathogen genes using TAC used to pre-screen samples for Context-Seq. Source data are provided in the Source Data file.

**Supplementary Table 2:** Taqman array card results for TEM and CTX-M in the 4 selected households. 1 indicates a positive detection (CT<40). Sex is indicated for sequenced human samples.

| HH | Sample | TEM | CTX-M Group 1 | CTX-M Group 9 | Selected? | Sex (M/F) |
|---|---|---|---|---|---|---|
| 1 | Older child | 1 | | 1 | | |
| 1 | Child | 1 | 1 | 1 | Y | F |
| 1 | Poultry | 1 | 1 | 1 | Y | |
| 1 | Dog | 1 | 1 | 1 | Y | |
| 2 | Older child | 1 | 1 | 1 | | |
| 2 | Adult | 1 | 1 | 1 | Y | F |
| 2 | Child | 1 | 1 | | | |
| 2 | Dog | 1 | 1 | 1 | Y | |
| 2 | Poultry 1 | 1 | 1 | 1 | | |
| 2 | Poultry 2 | 1 | 1 | 1 | | |
| 2 | Poultry 3 | 1 | 1 | 1 | Y | |
| 4 | Adult | 1 | 1 | 1 | Y | F |
| 4 | Child | 1 | 1 | | | |
| 4 | Poultry 1 | 1 | 1 | 1 | Y | |
| 4 | Poultry 2 | 1 | 1 | 1 | | |
| 4 | Poultry 3 | 1 | | | | |
| 3 | Older child | 1 | 1 | | Y | M |
| 3 | Adult | 1 | 1 | | | |
| 3 | Child | 1 | 1 | | Y | M |
| 3 | Poultry 1 | 1 | 1 | 1 | Y | |
| 3 | Dog | 1 | 1 | 1 | Y | |

**Supplementary Table 3**: Candidate guide RNA screened for use in this study. Mean and standard deviation of off-target hits and off-target scores were calculated by running the custom off-target script on seven human fecal samples from rural Kenya (PRJNA768833).[7]

| Guide sequence (5'-3') | Name | Strand | Genomic Location (nt) | Efficiency % | N Off-Target Hits Mean (std) | Off-Target Score Mean (std) |
|---|---|---|---|---|---|---|
| **TEM** | | | | | | |
| AGATCAGTTGGGTGCACGAGTGG | BT[1] | + | 105 | 54.47 | 480 (250) | 6 (3) |
| ACCGCGAGACCCACGCTCACCGG | BR | - | 704 | 51.04 | 339 (132) | 16 (9) |
| ATACGGGAGGGCTTACCATCTGG | CF[1] | - | 745 | 48.80 | 346 (97) | 33 (8) |
| ATCGAACTGGATCTCAACAGCGG | AI | + | 133 | 79.59 | 1308 (240) | 9 (6) |
| **CTX-M** | | | | | | |
| TGGCACCACCAACGATATCGCGG | CTX-1 | - | 729 | 69.45 | 2820 (417) | 18 (2) |
| CCAGTTCACGCTGATGGCGACGG | BV[1] | + | 21 | 59.2 | 3382 (534) | 24 (5) |
| GGTTTTATCCCCCACAACCCAGG | AQ[1] | - | 692 | 58.14 | 1188 (463) | 10 (2) |
| TTCACTTTTCTTCAGCACCGCGG | CTX-2 | - | 242 | 66.44 | 1019 (305) | 13 (3) |
| AAGTAAGTGACCAGAATCAGCGG | CTX-3 | - | 775 | 68.31 | 448 (71) | 21 (6) |
| GCCGCTGTATGCGCAAACGGCGG | CTX-4 | + | 72 | 66.56 | 1102 (613) | 16 (2) |

[1]Guide used in this study.

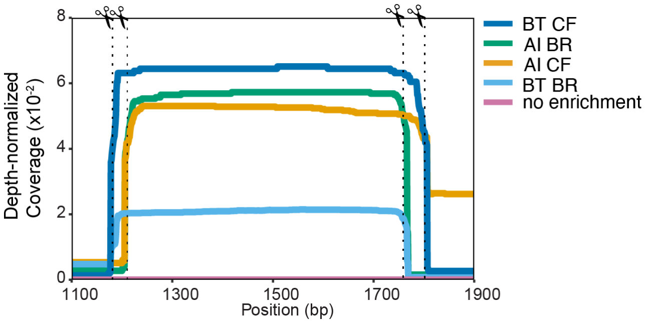


**Supplementary Figure 2**: Comparison between guide sets in **Supplementary Table 3** for TEM in our test system. Source data are provided in the Source Data file.

**Supplementary Table 4**: CTX-M alleles captured with CTX-M guides used in this study.

| Allele | BV | AQ | Allele | BV | AQ | Allele | BV | AQ |
|---|---|---|---|---|---|---|---|---|
| CTX-M-1 | | 1 | CTX-M-163 | 1 | 1 | CTX-M-88 | 1 | 1 |
| CTX-M-10 | 1 | 1 | CTX-M-164 | | 1 | CTX-M-96 | 1 | 1 |
| CTX-M-12 | 1 | 1 | CTX-M-166 | | 1 | CTX-M-101 | 1 | 1 |
| CTX-M-23 | | 1 | CTX-M-167 | 1 | 1 | CTX-M-103 | 1 | 1 |
| CTX-M-28 | | 1 | CTX-M-169 | 1 | 1 | CTX-M-114 | 1 | 1 |
| CTX-M-29 | 1 | 1 | CTX-M-170 | 1 | 1 | CTX-M-116 | | 1 |
| CTX-M-30 | | 1 | CTX-M-172 | 1 | 1 | CTX-M-117 | 1 | 1 |
| CTX-M-32 | | 1 | CTX-M-173 | 1 | 1 | CTX-M-123 | 1 | |
| CTX-M-33 | 1 | 1 | CTX-M-175 | | 1 | CTX-M-132 | 1 | 1 |
| CTX-M-34 | 1 | 1 | CTX-M-176 | 1 | 1 | CTX-M-136 | 1 | 1 |
| CTX-M-36 | | 1 | CTX-M-177 | 1 | 1 | CTX-M-139 | 1 | 1 |
| CTX-M-42 | 1 | 1 | CTX-M-179 | 1 | 1 | CTX-M-142 | 1 | 1 |
| CTX-M-53 | 1 | 1 | CTX-M-180 | 1 | 1 | CTX-M-150 | 1 | 1 |
| CTX-M-54 | 1 | 1 | CTX-M-181 | 1 | 1 | CTX-M-155 | 1 | 1 |
| CTX-M-58 | | 1 | CTX-M-182 | 1 | 1 | CTX-M-156 | 1 | 1 |
| CTX-M-60 | 1 | 1 | CTX-M-183 | 1 | 1 | CTX-M-157 | 1 | 1 |
| CTX-M-61 | | 1 | CTX-M-184 | 1 | 1 | CTX-M-158 | | 1 |
| CTX-M-62 | | 1 | CTX-M-186 | 1 | 1 | CTX-M-162 | 1 | 1 |
| CTX-M-64 | 1 | 1 | CTX-M-188 | 1 | 1 | CTX-M-37 | 1 | 1 |
| CTX-M-66 | 1 | 1 | CTX-M-189 | 1 | 1 | CTX-M-22 | 1 | 1 |
| CTX-M-68 | 1 | 1 | CTX-M-190 | 1 | 1 | CTX-M-15 | 1 | 1 |
| CTX-M-69 | 1 | 1 | CTX-M-193 | 1 | 1 | CTX-M-127 | 1 | 1 |
| CTX-M-71 | 1 | 1 | CTX-M-194 | 1 | 1 | CTX-M-137 | | 1 |
| CTX-M-72 | 1 | 1 | CTX-M-197 | 1 | 1 | CTX-M-138 | | 1 |
| CTX-M-80 | 1 | 1 | CTX-M-199 | 1 | 1 | CTX-M-144 | 1 | 1 |
| CTX-M-82 | 1 | 1 | CTX-M-202 | 1 | 1 | CTX-M-3 | 1 | 1 |
| CTX-M-209 | 1 | 1 | CTX-M-203 | 1 | | CTX-M-79 | 1 | 1 |
| CTX-M-210 | 1 | 1 | CTX-M-204 | 1 | 1 | CTX-M-55 | 1 | 1 |
| CTX-M-211 | 1 | 1 | CTX-M-206 | | 1 | CTX-M-52 | 1 | 1 |
| CTX-M-212 | 1 | 1 | CTX-M-207 | 1 | 1 | CTX-M-218 | 1 | 1 |
| CTX-M-216 | 1 | 1 | CTX-M-208 | 1 | 1 | CTX-M-220 | 1 | 1 |

**Supplementary Table 5**: TEM alleles captured with TEM guides used in this study

| Allele | BT | CF | Allele | BT | CF | Allele | BT | CF | Allele | BT | CF |
|---|---|---|---|---|---|---|---|---|---|---|---|
| TEM-1A | 1 | 1 | TEM-86 | 1 | 1 | TEM-149 | 1 | 1 | TEM-169 | 1 | 1 |
| TEM-1B | 1 | 1 | TEM-87 | 1 | 1 | TEM-150 | 1 | 1 | TEM-144 | 1 | 1 |
| TEM-1C | 1 | 1 | TEM-88 | 1 | 1 | TEM-151 | 1 | 1 | TEM-129 |  | 1 |
| TEM-1D | 1 | 1 | TEM-90 | 1 | 1 | TEM-152 | 1 | 1 | TEM-128 | 1 | 1 |
| TEM-2 |  | 1 | TEM-91 | 1 | 1 | TEM-153 | 1 | 1 | TEM-181 | 1 | 1 |
| TEM-2 |  | 1 | TEM-92 | 1 | 1 | TEM-154 | 1 | 1 | TEM-191 | 1 | 1 |
| TEM-3 |  | 1 | TEM-93 | 1 | 1 | TEM-155 |  | 1 | TEM-196 | 1 | 1 |
| TEM-6 | 1 | 1 | TEM-94 | 1 | 1 | TEM-156 | 1 | 1 | TEM-210 | 1 | 1 |
| TEM-8 |  | 1 | TEM-95 | 1 | 1 | TEM-157 | 1 | 1 | TEM-212 | 1 | 1 |
| TEM-10 | 1 | 1 | TEM-96 | 1 | 1 | TEM-158 | 1 | 1 | TEM-213 | 1 | 1 |
| TEM-11 |  | 1 | TEM-97 | 1 | 1 | TEM-159 | 1 | 1 | TEM-214 | 1 | 1 |
| TEM-12 | 1 | 1 | TEM-98 | 1 | 1 | TEM-160 |  | 1 | TEM-215 | 1 | 1 |
| TEM-15 | 1 | 1 | TEM-99 | 1 | 1 | TEM-162 |  | 1 | TEM-216 |  | 1 |
| TEM-16 |  | 1 | TEM-101 |  | 1 | TEM-163 | 1 | 1 | TEM-217 | 1 | 1 |
| TEM-17 | 1 | 1 | TEM-102 | 1 | 1 | TEM-164 |  | 1 | TEM-219 | 1 | 1 |
| TEM-20 | 1 | 1 | TEM-104 | 1 | 1 | TEM-166 | 1 | 1 | TEM-220 | 1 | 1 |
| TEM-21 |  | 1 | TEM-105 | 1 | 1 | TEM-167 | 1 | 1 | TEM-224 | 1 | 1 |
| TEM-22 |  | 1 | TEM-106 | 1 | 1 | TEM-168 | 1 | 1 | TEM-225 | 1 | 1 |
| TEM-24 |  | 1 | TEM-107 | 1 | 1 | TEM-171 | 1 | 1 | TEM-226 | 1 | 1 |
| TEM-28 | 1 | 1 | TEM-108 | 1 | 1 | TEM-176 | 1 | 1 | TEM-227 |  | 1 |
| TEM-29 | 1 | 1 | TEM-109 | 1 | 1 | TEM-177 |  | 1 | TEM-229 | 1 |  |
| TEM-30 | 1 | 1 | TEM-110 | 1 | 1 | TEM-178 |  | 1 | TEM-230 | 1 | 1 |
| TEM-33 | 1 | 1 | TEM-111 | 1 | 1 | TEM-182 | 1 | 1 | TEM-231 | 1 | 1 |
| TEM-34 | 1 | 1 | TEM-112 | 1 | 1 | TEM-183 | 1 | 1 | TEM-232 | 1 | 1 |
| TEM-43 | 1 | 1 | TEM-113 |  | 1 | TEM-184 | 1 | 1 | TEM-233 | 1 | 1 |
| TEM-45 | 1 | 1 | TEM-114 |  | 1 | TEM-185 | 1 | 1 | TEM-234 | 1 | 1 |
| TEM-47 | 1 | 1 | TEM-115 | 1 | 1 | TEM-186 | 1 | 1 | TEM-148 | 1 | 1 |
| TEM-48 | 1 | 1 | TEM-116 | 1 | 1 | TEM-187 | 1 | 1 | TEM-40 | 1 | 1 |
| TEM-49 | 1 | 1 | TEM-120 | 1 | 1 | TEM-188 | 1 | 1 | TEM-36 | 1 | 1 |
| TEM-52C | 1 | 1 | TEM-121 |  | 1 | TEM-189 | 1 | 1 | TEM-35 | 1 | 1 |
| TEM-52B | 1 | 1 | TEM-122 | 1 | 1 | TEM-190 | 1 | 1 | TEM-32 | 1 | 1 |
| TEM-53 | 1 | 1 | TEM-123 | 1 | 1 | TEM-193 | 1 | 1 | TEM-26 | 1 | 1 |
| TEM-54 | 1 | 1 | TEM-124 | 1 | 1 | TEM-194 | 1 | 1 | TEM-19 | 1 | 1 |
| TEM-55 | 1 | 1 | TEM-125 | 1 | 1 | TEM-195 | 1 | 1 | TEM-9 | 1 | 1 |
| TEM-57 | 1 | 1 | TEM-126 | 1 | 1 | TEM-197 | 1 | 1 | TEM-4 | 1 | 1 |
| TEM-60 |  | 1 | TEM-127 | 1 | 1 | TEM-198 | 1 | 1 | TEM-138 | 1 | 1 |
| TEM-63 | 1 | 1 | TEM-130 |  | 1 | TEM-201 | 1 | 1 | TEM-139 |  | 1 |
| TEM-67 |  | 1 | TEM-131 | 1 | 1 | TEM-205 | 1 | 1 | TEM-141 | 1 | 1 |
| TEM-68 | 1 | 1 | TEM-132 | 1 | 1 | TEM-206 | 1 | 1 | TEM-142 | 1 | 1 |
| TEM-70 | 1 | 1 | TEM-133 | 1 | 1 | TEM-207 | 1 | 1 | TEM-143 | 1 | 1 |
| TEM-71 | 1 | 1 | TEM-134 |  | 1 | TEM-208 | 1 | 1 | TEM-145 | 1 | 1 |
| TEM-72 |  | 1 | TEM-135 | 1 | 1 | TEM-209 |  | 1 | TEM-146 | 1 | 1 |
| TEM-76 | 1 | 1 | TEM-136 | 1 | 1 | TEM-211 | 1 | 1 | TEM-147 | 1 | 1 |
| TEM-77 | 1 | 1 | TEM-137 | 1 | 1 | TEM-52 | 1 | 1 | TEM-84 | 1 | 1 |
| TEM-78 | 1 | 1 | TEM-80 | 1 | 1 | TEM-82 | 1 | 1 | TEM-85 | 1 | 1 |
| TEM-79 | 1 | 1 | TEM-81 | 1 | 1 | TEM-83 | 1 | 1 |  |  |  |

**Supplementary Table 6:** MinION flow cell ID numbers and SRA BioSample numbers for BioProject PRJNA1157857.

| Sample | Flow Cell Number | SRA BioSample |
|---|---|---|
| **HH samples** | | |
| 1_child_feces | FAW10902 | SAMN43524419 |
| 1_poultry_feces | FAW10718<br>FAW05518 | SAMN43524420 |
| 1_dog_feces | FAW10742 | SAMN43524421 |
| 2_adult_feces | FAS08862 | SAMN43524422 |
| 2_dog_feces | FAS96515 | SAMN43524423 |
| 2_poultry_feces | FAX19523 | SAMN43524424 |
| 3_child_feces | FAS07765 | SAMN43524425 |
| 3_older_child_feces | FAS08862 | SAMN43524426 |
| 3_dog_feces | FAS10189 | SAMN43524427 |
| 3_poultry_feces | FAS08528 | SAMN43524428 |
| 4_adult_feces | FAS97950 | SAMN43524429 |
| 4_poultry1_feces | FAS08896 | SAMN43524430 |
| 4_poultry2_feces | FAS40845 | SAMN43524431 |
| **Optimization** | | |
| regular | FAP82268 | SAMN43584091 |
| adaptive | FAQ52471 | SAMN43584092 |
| 2 guides | FAQ49828 | SAMN43584093 |
| 2 hr cas9 | FAP85007 | SAMN43584094 |
| 2 targets | FAQ49794 | SAMN43584095 |
| no enrichment | FAQ52008 | SAMN43584096 |
| ProK | FAS96541 | SAMN43584097 |
| Regular enrichment (for proK comparison) | FAS08270 | SAMN43584098 |
| CTX-M only | FAS09073 | SAMN43584099 |
| TEM only | FAS09238 | SAMN43584100 |
| CTX-M and tem together | FAS07765 | SAMN43584101 |
| **TEM Guides[1]** | | |
| TEM_AI_BR | AJN275 | SAMN43584102 |
| TEM_AI_CF | AJM918 | SAMN43584103 |
| TEM_BT_BR | AJN252 | SAMN43584104 |
| TEM_CF_BT | AJM933 | SAMN43584105 |
| TEM_control | AJN279 | SAMN43584106 |

[1]Flongle flow cell

**Supplementary Table 7:** Sequencing summary for samples used in this work. Total number of reads, reads containing ARGs, average length of reads, and number of clusters used to generate consensus sequences.

| | | | TEM | | | CTX-M | | |
|---|---|---|---|---|---|---|---|---|
| HH | Sample | Total # of reads | # of reads | Ave length | Number of clusters | # of reads | Ave length | Number of clusters |
| 1 | Child feces | 227,975 | 79 | 4314 | 10 | 0 | - | - |
| 1 | Poultry feces | 266,102 | 443 | 4509 | 30 | 0 | - | - |
| 1 | Dog feces | 127,865 | 856 | 5886 | 58 | 30 | 3921 | 6 |
| 2 | Adult feces | 2,669,501 | 1398 | 4956 | 15 | 0 | - | - |
| 2 | Dog feces | 1,909,047 | 3235 | 1818 | 50 | 419 | 1556 | 10 |
| 2 | Poultry feces | 148,896 | 2948 | 4876 | 116 | 47 | 4741 | 6 |
| 3 | Child feces | 339,046 | 1778 | 5268 | 28 | 68 | 5508 | 5 |
| 3 | Older child feces | 993,679 | 223 | 4858 | 15 | 3 | 7106 | 0 |
| 3 | Dog Feces | 1,915,359 | 6768 | 6162 | 86 | 0 | - | - |
| 3 | Poultry feces | 199,895 | 204 | 4286 | 27 | 33 | 3278 | 5 |
| 4 | Adult feces | 172,759 | 59 | 5751 | 3 | 48 | 4559 | 4 |
| 4 | Poultry 1 feces | 398,520 | 341 | 5385 | 12 | 0 | - | - |
| 4 | Poultry 2 feces | 109,002 | 1128 | 5037 | 22 | 0 | - | - |
